# Responses to unannounced rule changes in adults with developmental disabilities: A community-based pilot study with consumer-grade EEG

**DOI:** 10.64898/2026.10.17.759777

**Authors:** Yunjeong Jeong, Suho Hyun

## Abstract

Unannounced changes in rules or environments often precede challenging behavior and limit community participation for adults with developmental disabilities, yet their responses to change have rarely been quantified directly. We recruited adults with developmental disabilities, including those with intellectual disability, at a community day-service center. We administered a probabilistic reversal learning task and recorded resting-state electroencephalography (EEG) via a 14-channel wireless EEG system. Among the 12 adults who performed the task, total errors were comparable across participants (range 31–57), whereas a switch bias index, contrasting perseverative errors (persisting with the previously correct option) with regressive errors (failing to maintain the newly correct option), ranged from −0.96 to +0.91. In the 9 participants included in resting-state EEG analyses, a flatter aperiodic spectrum over the frontal cluster (preregistered, but selected as the focus post hoc) was associated with fewer switches after negative feedback (ρ = .80) and more perseverative errors (ρ = −.72). The preregistered parieto-occipital cluster showed the same direction (ρ = .69, 95% confidence interval [−.11, .97], *n* = 8), which was opposite to the preregistered prediction. Frontal alpha peak amplitude was associated with accuracy (ρ = .77), and the between-run reliability of resting-state measures was .82–.99. These findings provide preliminary evidence that EEG recorded in community settings can capture individual differences in how people respond to change. Field measurement that includes people with intellectual disability may help extend the evidence base for the Sustainable Development Goals of good health and well-being (SDG 3) and reduced inequalities (SDG 10).

## 1. Introduction

For adults with developmental disabilities, unannounced change is a routine part of community life, such as a shifted schedule, an altered rule or an unfamiliar setting. In support settings, episodes of challenging behavior and difficulty joining others often coincide with such changes. Challenging behavior increases when events are unpredictable (Flannery & Horner, 1994), whereas previewing the sequence of activities reduces challenging behavior (Lequia et al., 2012). In adults with intellectual disability and autistic adults, lower set-shifting scores have been associated with aggression (Visser et al., 2014). Yet when faced with the same change, some people adjust quickly while others persist with the previous routine. This variability is frequently noted in practice, but what gives rise to it remains poorly understood.

The question extends beyond individual behavior. For people who find change difficult, unfamiliar settings and altered rules become barriers to participation. The challenging behavior that follows is often recorded as a problem of the individual, which in turn distances them from community activities. Understanding how individuals differ in responses to change would enable support that, rather than eliminating change, anticipates change and supports each person through it in ways that suit them. It thus bears directly on two Sustainable Development Goals (SDGs): good health and well-being (SDG 3) and reduced inequalities (SDG 10).

### 1.1 Cognitive flexibility and reversal learning

Cognitive flexibility is understood as the ability to adjust behavior in response to changing circumstances (Uddin, 2021). After a change, this adjustment can be decomposed into two steps: disengaging from the previously appropriate response and maintaining the newly appropriate one. Difficulty with the first step appears as perseverative errors, that is, continuing to choose the previously correct option; instability in the second step appears as regressive errors (a task term unrelated to developmental regression), that is, returning to the previous option after the new one has been chosen. Probabilistic reversal learning tasks dissociate these two error types within a single task and have been widely used in research on autism and developmental disabilities (Crawley et al., 2020; D’Cruz et al., 2013).

### 1.2 Why individual differences

Variation in cognitive flexibility may be better captured at the individual level than by group means. Reviewing 72 studies, Leung and Zakzanis (2014) found that no performance-based measure reliably distinguished autistic from non-autistic people. In a meta-analysis of 59 studies, Lage et al. (2024) estimated small-to-moderate group differences with substantial heterogeneity, the largest effect being on perseverative errors. Individual studies also diverged in direction: D’Cruz et al. (2013) reported more regressive errors in autistic participants, whereas Crawley et al. (2020) reported a stronger perseverative tendency.

Perseverative and regressive errors reflect opposite tendencies. Total error scores conflate the two, and when individuals lean in opposite directions, group means may cancel these tendencies. Individual distributions of error types may therefore be more informative than group-level mean error counts. Accordingly, we describe the distribution of error types before testing hypotheses.

### 1.3 Including people with intellectual disability

Samples in cognitive flexibility research typically exclude participants with intellectual disability in order to dissociate task performance from intellectual functioning (Crawley et al., 2020; Lage et al., 2024). Because more than one third of autistic children also have intellectual disability (37.9%; Maenner et al., 2023), it remains unknown whether such findings apply to them. From the perspective of SDG 10, which calls for the social inclusion of all irrespective of disability, this exclusion represents both a gap in the evidence and an imbalance in who participates in research.

Restricting samples by intellectual functioning may also remove the variation of interest. In adults with mild or borderline intellectual disability, an autism diagnosis did not predict shifting scores (Roelofs et al., 2015), and aggression was associated with shifting scores rather than with diagnosis (Visser et al., 2014). Individual differences in shifting thus exist independently of diagnostic categories. We therefore did not restrict our sample by diagnosis or intellectual level.

### 1.4 Measurement constraints

Measuring responses to changing settings in this population has faced two constraints. First, widely used self-report scales have not been validated in this population. Second, neuroimaging is difficult to apply: magnetic resonance imaging requires lying still in a confined space for a long time, and sedation, which is often used to make this possible, may confound the measurements of brain activity (Smith et al., 2019; Stogiannos et al., 2022); even without sedation, autistic children are more often excluded for head motion (Nebel et al., 2022).

Mobile EEG has been proposed as a way around these constraints (Lau-Zhu et al., 2019), as it provides a physiological measure that does not depend on self-report. With few electrodes, no conductive gel and the possibility of recording on site, it places little burden on participants. Its validation, however, has mostly been carried out in nondisabled adults, and which measures remain reliable in field settings with frequent poor electrode–skin contact and movement must be checked separately.

### 1.5 The present study

We administered a probabilistic reversal learning task and recorded resting-state EEG in adults with developmental disabilities attending a community day-service center. Behaviorally, we separated perseverative and regressive errors to characterize whether individuals tended to stay with the previous option or to return to it after switching. For EEG, the primary measure was the aperiodic component of the power spectrum, which has been proposed as a non-invasive index of excitation– inhibition balance (Donoghue et al., 2020; Gao et al., 2017).

In lieu of group comparison, we examined individual differences within adults with developmental disabilities and addressed three questions:

1. *How much do responses after reversal differ between individuals?*
2. *Are these differences associated with resting-state EEG?*
3. *Does the EEG measure change within individuals immediately after reversal?*

The primary hypothesis was that the resting-state aperiodic exponent (primary cluster: parieto-occipital) would be associated with the switch bias index. Within-person hypotheses predicted that the exponent would be lower in the eight trials after a reversal than in the eight trials before it (W1, parieto-occipital) and that post-feedback theta would be larger after unrewarded than after rewarded trials (W2, frontal). Perseverative and regressive error counts were co-primary outcomes, as was the reinforcement-learning parameter κ, which could not be identified in these data and was replaced by the lose-shift rate (Table S1); alpha-band activity and informant-reported sameness scores on the Adult Repetitive Behaviours Questionnaire-2 (RBQ-2A; Barrett et al., 2015, 2018; Brett et al., 2024) were exploratory. As a pilot study, the present investigation aimed to assess feasibility and estimate effect sizes.

## 2. Materials and methods

### 2.1 Ethics and preregistration

The study was conducted with the cooperation of a community center providing daytime activity services for adults with developmental disabilities (hereafter, the center). The study did not undergo formal review by an institutional review board (IRB; see Declarations). All participants were adults and provided written informed consent in person after the study had been explained to them.

The hypotheses and outcomes described in Section 1.5, together with the analysis plan, were preregistered on the Open Science Framework (OSF; https://osf.io/qvb9p; registered August 22, 2026). Registration took place after data from the first 10 participants had been collected; before registration, data quality and descriptive behavioral statistics had been examined for the first five participants, but no EEG–behavior associations had been computed. Deviations from the preregistration are listed in Table S1 and described in Supplementary Appendix 1.

### 2.2 Participants

Participants were recruited through notices posted at the center. Eligibility required a registered developmental disability and the ability to attend to a screen and operate a computer mouse. Individuals who did not pass the practice block were excluded. By protocol, individuals under adult guardianship were also ineligible since consent was to be given by participants themselves; however, none of those approached were under guardianship. Co-occurring disabilities and intellectual level were not used as exclusion criteria. Sessions were conducted at the center and lasted approximately 20 min; no sedation or study-related medication was administered (Figure 2).

Although the preregistration and protocol specified autistic adults, eligibility in practice was based on a registered developmental disability. Of the 15 participants who completed the session, 4 had autism as their registered disability (Table 1). The results therefore cannot be interpreted as tests of hypotheses about autistic adults; this deviation is documented in Supplementary Appendix 1.

**Table 1.** Participant characteristics.

| Characteristic |  | Value |
| --- | --- | --- |
| Analyzed | — | 15 |
| Age (years) | Mean (SD) · range | 30.3 (10.5) · 21–59 |
| Sex | Male / female | 11 / 4 |
| Registered disability<sup>a</sup> | Intellectual disability | 9 |
|  | Autism | 4 |
|  | Developmental disability | 1 |
|  | Multiple (hearing, intellectual) | 1 |
| Conditions noted in registration | Down syndrome · brain lesion | 2 · 1 |
| Psychotropic medication | Yes / no | 9 / 6 |

Note. Demographics exclude the participant who withdrew. <sup>a</sup>Primary disability on the welfare card; Down syndrome and brain lesion appear within the intellectual disability registration and are included among the nine. Groups were too small for comparisons by type.

### 2.3 Task

On each trial of the probabilistic reversal learning task (Figure 1), a blue circle and an orange square were presented on the left and right of the screen, and participants selected one with the mouse. Choices of the correct option were rewarded (green O) with 80% probability; unrewarded correct choices and all incorrect choices were followed by no reward (red X; 80/0 schedule). Each trial consisted of fixation (0.4–0.7 s), stimulus presentation (up to 4 s), choice highlight (0.2 s), feedback (0.7 s) and an inter-trial interval (0.5 s).

**Figure 1.**
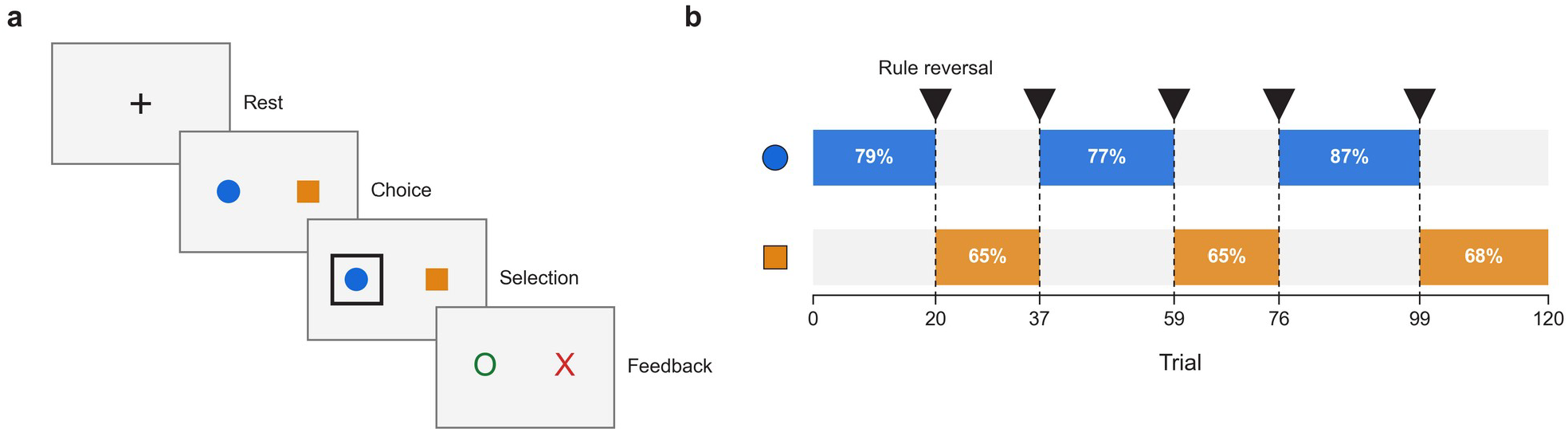
Task structure. **a**, Resting-state recording (eyes closed, two runs of 90 s) and the sequence of one trial. The chosen stimulus is outlined, followed by reward (O) or no reward (X); the left–right positions of the two stimuli vary across trials. **b**, Reversal schedule. Shaded segments mark the correct stimulus; numbers give the proportion of correct choices actually rewarded (incorrect choices, 0%). Dashed lines mark reversals; the schedule was identical for all participants

The task comprised 120 trials (three blocks of 40 without breaks), and the correct option reversed without warning five times (segments of 17–23 trials). The reward schedule was generated with a fixed seed and was identical for all participants; 89 of the 120 trials (74.2%) were scheduled for reward. Two practice stages preceded the task. To pass the first stage, participants had to choose the correct option on seven consecutive trials within 24 trials in which correct choices were always rewarded; in the second, they completed 12 trials under task conditions (one reversal) without a pass criterion.

Feedback-based strategy was characterized by lose-shift, win-stay and switch rates (Equation 1). These rates were computed over adjacent pairs of valid choices after non-responses were excluded, and accuracy was likewise computed over valid choices.

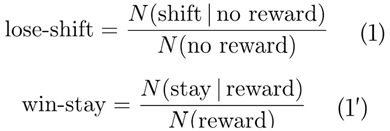

Errors after each reversal were classified by the task program as perseverative errors (*N*_pers_), made before the newly correct option was first chosen, or regressive errors (*N*_reg_), made thereafter. The balance between the two was summarized as the switch bias index (SBI; Equation 2), with values near +1 indicating predominantly regressive errors and values near −1 predominantly perseverative errors.

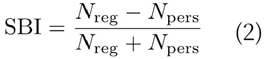

For each post-reversal segment, trials-to-recover was defined as the zero-based trial index at which the participant first completed three consecutive selections of the newly correct option; if this criterion was not met, the segment length was used. The mean across the five post-reversal segments served as each participant’s value, so the measure reflects both whether and how quickly the new rule was acquired. As preregistered, the effectiveness of the reversal manipulation was assessed as the proportion of choices of the newly correct option 6–15 trials after each reversal; this proportion was reported descriptively, without a threshold, and was not used for exclusion.

### 2.4 EEG recording and preprocessing

#### Recording

EEG was recorded with a consumer-grade wireless headset (Emotiv EPOC X; 14 channels, 128 Hz, hardware band-pass 0.2–45 Hz, fixed P3/P4 reference). The same headset has been used to record resting-state EEG from children in classrooms, where its aperiodic measures showed reliability comparable to that of a research-grade system (van Bueren et al., 2026). Two 90-s eyes-closed resting-state runs were recorded before the task, separated by a 20-s fixation screen and an instruction screen read with eyes open. Stimulus onsets were recorded as event markers, and the clock offset drift between the start and end of each session ranged from −0.70 to +0.68 ms, within the preregistered limit (20 ms).

#### Preprocessing

Data were processed with MNE-Python (Gramfort et al., 2013) and are reported following the COBIDAS-MEEG recommendations (Pernet et al., 2020). Bad channels were identified on a copy of the data filtered at 1–45 Hz with a 60-Hz notch, as channels whose median peak-to-peak amplitude exceeded three times, or fell below 0.2 times, the median across channels; these channels were excluded without interpolation. Only a 0.5-Hz high-pass filter was applied to the analyzed signal, because low-pass filtering attenuates high-frequency power and thereby inflates the aperiodic exponent, and the hardware already limits the bandwidth to 45 Hz. After the first 10 s of each run were discarded, the data were divided into non-overlapping 2-s segments; segments exceeding 200 µV peak-to-peak were rejected, and runs with less than 30 s of remaining data were excluded. Power spectra were computed with a Hann window and averaged in linear power (Welch, 1967), and periodic and aperiodic components were separated at 2–40 Hz with specparam (fixed mode; fits with R^2^ < .85 treated as missing; Donoghue et al., 2020). The aperiodic component is modeled by Equation 3, in which *χ* denotes the aperiodic exponent.

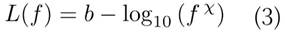

Independent component analysis was not applied, because ocular artifacts are minimal with eyes closed and, without electrooculography channels, components could not be classified objectively.

### 2.5 Analysis and statistics

The resting-state measures were the aperiodic exponent and alpha peak amplitude (7–14 Hz), computed for the four preregistered clusters: parieto-occipital (O1, O2, P7, P8), temporo-parietal (T7, T8, P7, P8), frontal (AF3, AF4, F3, F4) and global (all channels passing quality criteria). The frontal cluster was chosen as the focus of the correlational analyses after inspection of the data. The definition of alpha peak amplitude differs from the preregistered definition of alpha power (Table S1). Unless otherwise stated, EEG values are from the manuscript specification (original reference, 200-µV rejection threshold, R^2^ ≥ .85, at least 30 s per run); values under the preregistered specification (common average reference, 150 µV, R^2^ ≥ .90, at least 60 s per run, clusterwise epoch rejection) are reported alongside where relevant. Relative band power was expressed as a proportion of total 1–45 Hz power (delta 2–4, theta 4–8, alpha 8–13, beta 13–30, gamma 30–45 Hz).

Connectivity during the task was quantified as the alpha-band (8–12 Hz) phase-locking value (PLV; Lachaux et al., 1999), computed from the instantaneous phases *φ* of electrodes *j* and *k* as in Equation 4, where ⟨·⟩_*t*_ denotes the time average.

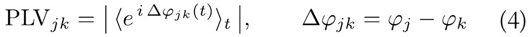

Pairwise values were averaged within regions, and characteristic path length (distance = 1/PLV) and clustering coefficient were computed on the weighted graph (Onnela et al., 2005). Because scalp PLV is affected by volume conduction, it was interpreted only as sensor-level synchrony.

Between-participant associations (preregistered for estimation only) were assessed with Spearman correlations. Rest–task comparisons are reported as paired differences (task − rest), together with the number of participants changing in the same direction (*k*/*n*). Intervals for correlations, partial correlations and differences between correlations (Δρ) are participant-level bootstrap 95% intervals (10,000 resamples, seed 1; resamples with fewer than four distinct participants discarded). Between-run reliability was quantified with intraclass correlation coefficients (ICCs; Koo & Li, 2016). The within-person hypotheses (W1, W2) were estimated with the preregistered linear mixed models and are reported with Wald 95% intervals; intervals for the three-level W2 contrasts were obtained with a participant cluster bootstrap.

Interpretation is based on estimates and intervals, with *P* values reported for reference (tests are listed in Table S4). The three preregistered tests (H1, W1, W2) were not corrected for multiple comparisons; the remaining exploratory tests were corrected with the Benjamini– Hochberg procedure within three families (11 rest–task changes, 16 EEG–behavior correlations and 18 additional W1/W2 contrasts), and asterisks denote *q* values (^*^ *q* < .05, ^**^ *q* < .01). Topographies (Figure 5a) were corrected with a cluster-based permutation test using participant-level sign flips (10,000 permutations, seed 1; Maris & Oostenveld, 2007).

## 3. Results

### 3.1 Participants and data collection

The analyzed samples comprised 12 participants for behavior, 9 for resting-state EEG and 10 for task EEG (Figure 2, Table 1). Of the 16 participants recruited, 1 withdrew after the first resting-state run and 3 did not pass the practice block (pass rate 12/15 = 80.0%), leaving 12 who performed the task. Eleven completed all 120 trials, and one (009) stopped at trial 98 after four reversals. Nonresponses accounted for 45 of 1,418 trials, 23 of which came from a single participant.

Three participants were excluded from the resting-state EEG sample: 007, for whom only one of the two runs met the quality criterion (and whose reaction times also exceeded the preregistered limit; Table S3), and 008 and 009, who met the preregistered exclusion criterion (winstay < .60). The exclusion of 007 follows the channel-wide epoch rejection of the manuscript specification and might not occur under the preregistered cluster-wise rule (Supplementary Appendix 1). Task EEG was analyzed in the 10 participants who met the artifact criterion for the task period; the two EEG samples therefore differ in composition (Table S2).

**Figure 2.**
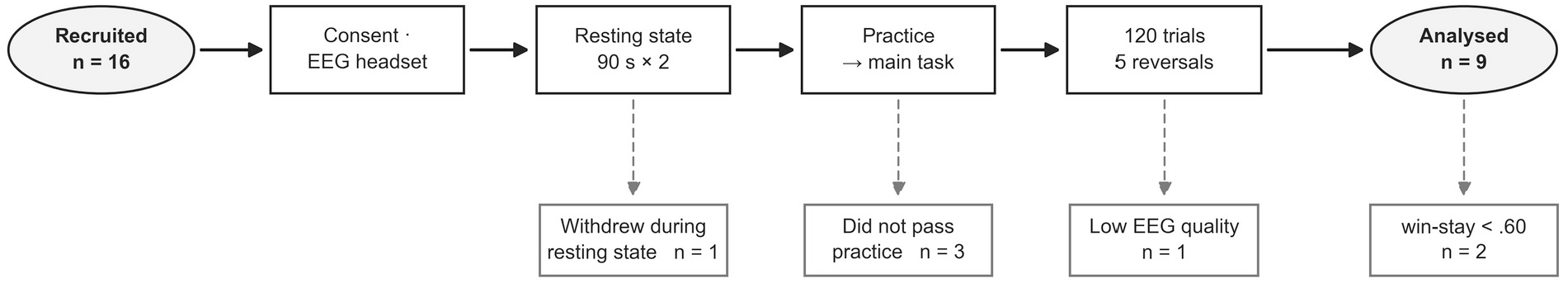
Session procedure and participant flow. The top row shows the order of procedures; boxes below give exclusions at each stage (EEG quality, 007; win-stay criterion, 008 and 009). Task EEG was analyzed in the 10 participants who met the artifact criterion for the task period

Behavioral analyses included all 12 participants who performed the task, in order to describe the full distribution; values for the 9 participants who remained after all preregistered exclusion criteria were applied (i.e., excluding 007, 008 and 009) are reported in Supplementary Appendix 1.

The reversal manipulation largely worked as intended. The proportion of choices of the newly correct option 6–15 trials after each reversal averaged 0.563 (*SD* = 0.094, range 0.380–0.720; for 009, the mean across four reversals) and exceeded 0.5 in 10 of 12 participants. The lowest value was observed for 004, who chose almost exclusively one shape (Section 3.2).

### 3.2 Error types and individual differences

Participants made comparable numbers of errors, but the composition of their errors diverged in opposite directions (Figure 3). Whereas total errors were relatively uniform (*M* = 40.2, *SD* = 7.5, range 31–57), the switch bias index ranged from −0.965 to +0.909 (*M* = +0.510). Perseverative-dominant and regressive-dominant participants thus coexisted within one group, and the two error counts were strongly negatively correlated (ρ = −.863). Two participants were perseverative-dominant, 004 (57 errors) and 005 (45 errors), and the 10 regressive-dominant participants made a median of 36 errors. Because only two participants fell on one side, individual values are presented instead of group comparisons. The switch bias index was defined for all participants, each of whom made at least three perseverative and regressive errors combined.

**Figure 3.**
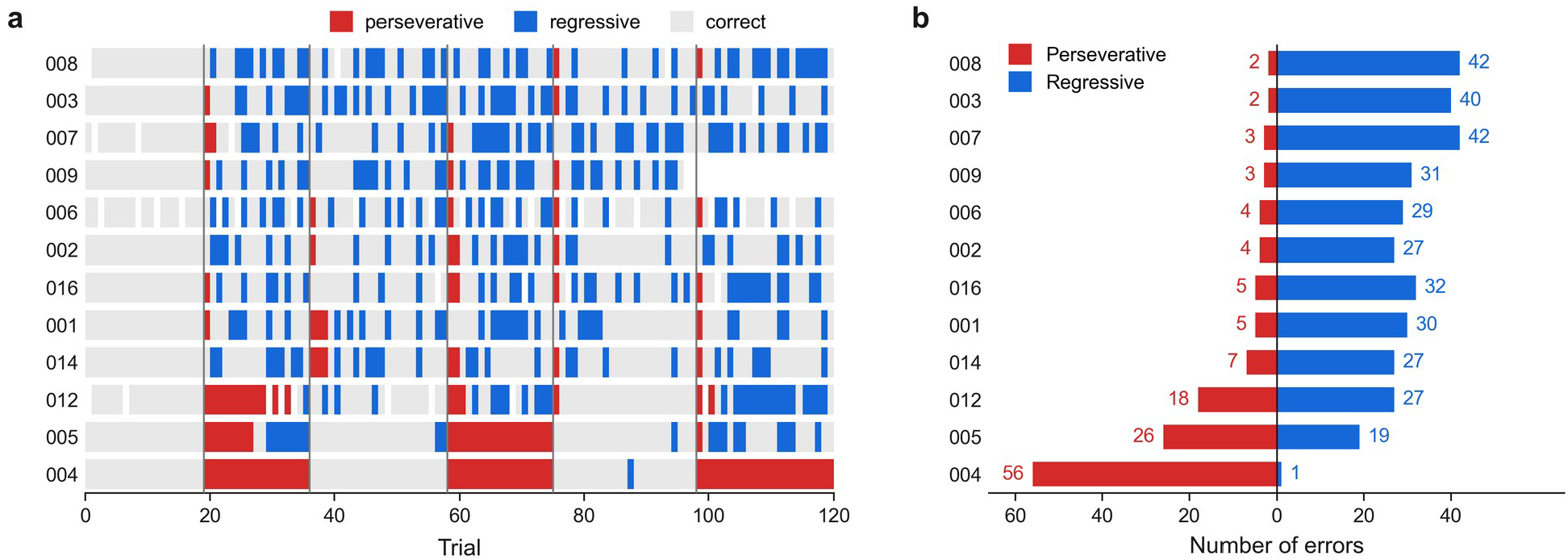
Distribution of error types. **a**, Trial records by participant (one row per participant, one cell per trial). Red, perseverative errors; blue, regressive errors; gray, correct choices and acquisition errors in the first segment; blank, non-responses. Vertical lines mark reversals; participants are ordered by switch bias index. **b**, Error counts by participant

The two most extreme participants illustrate this divergence. Participant 004 chose the same shape on 118 of 120 trials, maintaining it through four reversals during the longest run (82 trials). By contrast, 003 switched on about every second trial (switch rate 50.8%, lose-shift 62.3%). Their total errors were similar (57 and 42), showing that summed error scores do not capture this difference.

For the other behavioral measures, means were 11.3 perseverative errors (*SD* = 15.9), 28.9 regressive errors (*SD* = 11.2), a lose-shift rate of .508, a win-stay rate of .765 and an accuracy of .597 (range .517–.700). The obtained reward rate (*M* = .435) was lower than the maximum attainable under the fixed schedule (.742).

### 3.3 Resting-state EEG and behavior

Two frontal resting-state measures were associated with distinct aspects of post-reversal behavior (Figure 4d– h): the aperiodic exponent to the direction of responses, and alpha peak amplitude to the level of performance. The bootstrap 95% confidence intervals excluded zero for all five correlations.

**Figure 4.**
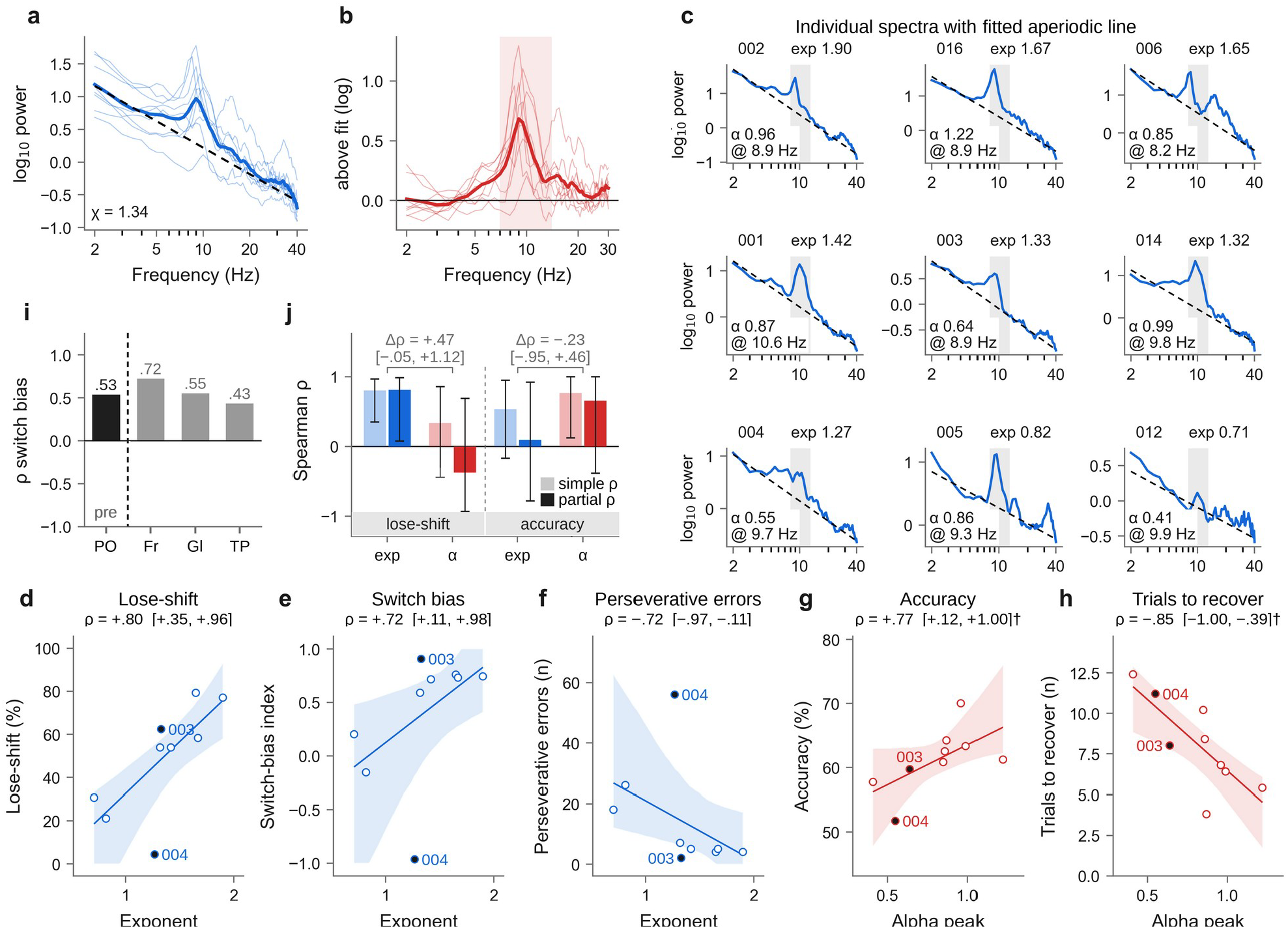
Resting-state EEG and behavior. **a**, Frontal spectra with aperiodic fits (dashed); thin lines are individual participants. **b**, Spectra with the aperiodic component removed; shading marks the alpha peak search range (7–14 Hz). **c**, Frontal spectra by participant (title, participant and exponent; inset, alpha peak amplitude and center frequency). **d–h**, Associations between resting-state and behavioral measures (*n* = 9). Filled points are the cases described in the text (003, 004); ρ is the Spearman correlation with its bootstrap 95% interval, and † indicates an interval reaching ±1.00. Fit lines and shading are for visualization (Table S1). **i**, Correlation of aperiodic exponent with switch bias index by cluster (PO, parieto-occipital; Fr, frontal; Gl, global; TP, temporo-parietal; PO and Fr, *n* = 9; Gl, 8; TP, 7); pre, preregistered primary cluster (manuscript specification). **j**, Simple and partial correlations (exp, aperiodic exponent; α, alpha peak amplitude) with bootstrap 95% intervals; Δρ, difference between the two EEG correlations

Participants with a smaller exponent (i.e., a flatter spectrum) switched less often after negative feedback (lose-shift ρ = .80 [.35, .96]), made more perseverative errors (ρ = −.72 [−.97, −.11]) and had a switch bias index shifted toward perseveration (ρ = .72 [.11, .98]; Figure 4d–f). These associations were robust to analytic choices: leaving out one participant at a time yielded .74–.84 for lose-shift, the lose-shift association held under alternative exclusion criteria (the association with the switch bias index weakened when no criteria were applied), and all 41 preregistered specifications that met the minimum sample size yielded positive estimates (Supplementary Appendix 1). Because the protocol predicted that a flatter spectrum would accompany greater flexibility, the observed direction was opposite to the prediction.

Participants with a larger alpha peak were more accurate (ρ = .77 [.12, 1.00]) and required fewer trials to reach the new rule after reversals (ρ = −.85 [−1.00, −.39]; Figure 4g–h). Alpha peak amplitude was not associated with measures of response direction, such as the switch bias index or perseverative errors, and this dissociation was maintained in partial correlations controlling for the other EEG measure (Figure 4j).

The association between the exponent and the switch bias index was positive in all four preregistered clusters (Figure 4i); in the preregistered primary cluster (parietooccipital) under the preregistered specification, it was ρ = .69 [−.11, .97] (*n* = 8). It should be noted, however, that the focus on the frontal cluster was chosen post hoc, that the analyses of accuracy, trials to recover and partial correlations were not preregistered, and that, with nine participants, the *q* value within the family of 16 EEG– behavior correlations was .077 (Table S4).

### 3.4 EEG during the task

Relative to rest, alpha activity and alpha synchrony both decreased during the task (Figure 5). All analyses in this section were exploratory; values in parentheses are paired differences (task − rest) with bootstrap 95% intervals.

**Figure 5.**
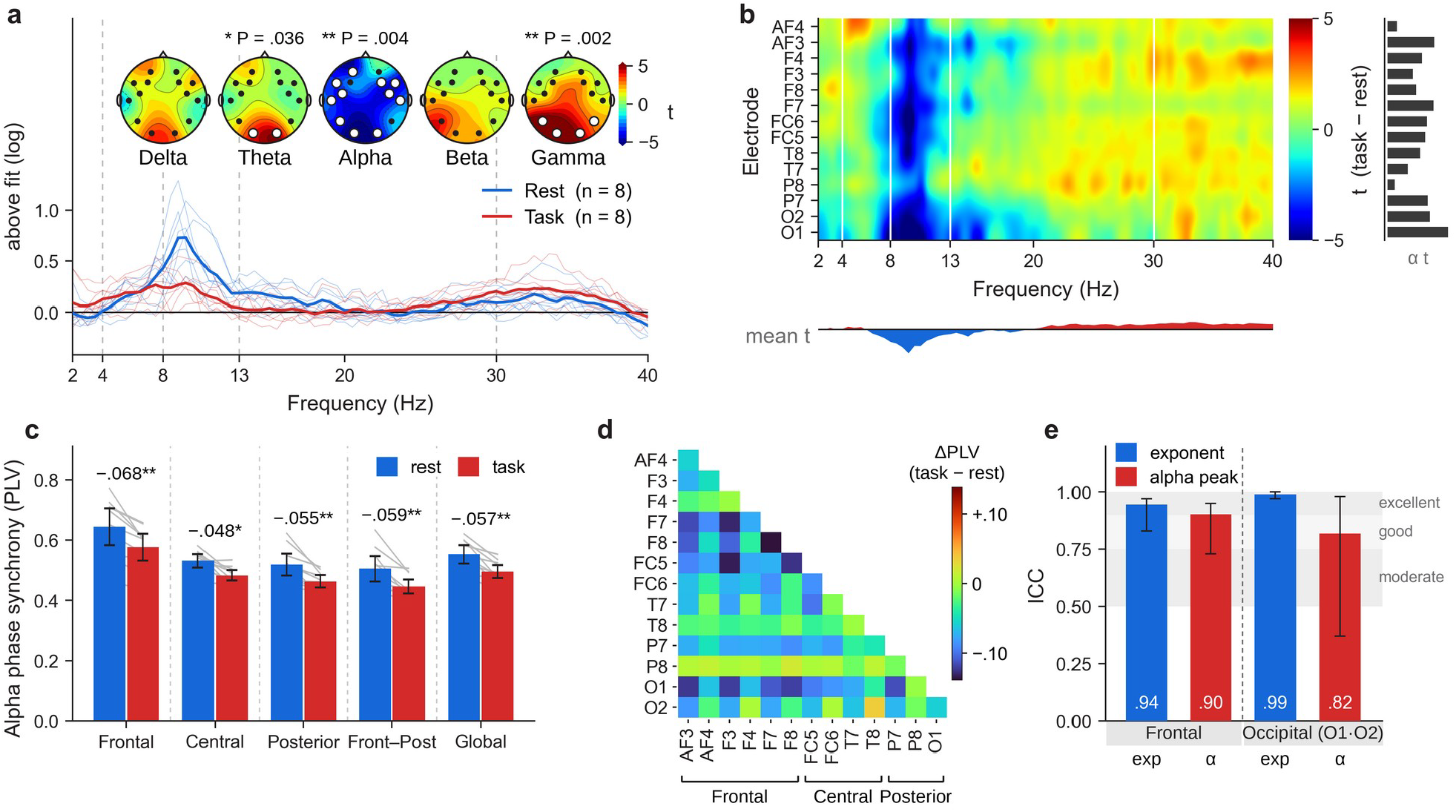
Change from rest to task. **a**, Frontal spectra with the aperiodic component removed (*n* = 8). Topographies show paired *t* for channel relative power by band (*n* = 10); *P* values above are cluster-permutation corrected, and white dots mark electrodes in surviving clusters. **b**, Paired *t* of log power by electrode (task − rest); right, mean alpha *t* per electrode; bottom, mean *t* per frequency; values beyond ±5 are shown in the end colors. **c**, Alpha phase-locking value (PLV) by region with 95% confidence intervals; numbers above bars are task − rest differences and asterisks family-corrected *q* values (^*^ *q* < .05, ^**^ *q* < .01). **d**, Change in alpha PLV by electrode pair (task − rest, mean of 10); brackets mark the regions in c; pairs were not tested individually. **e**, Intraclass correlations between the two resting-state runs with bootstrap 95% intervals (occipital, O1 and O2)

Frontal relative alpha power decreased from .227 to .124 (−0.102 [−0.156, −0.052], *q* < .05^*^, 9 of 10 participants), and periodic alpha (after removal of the aperiodic component) and alpha peak amplitude decreased in the same direction (Supplementary Appendix 3). Relative delta power increased (+0.042 [+0.015, +0.073], *q* < .05^*^), whereas the intervals for theta, beta and gamma included zero. Increases above 20 Hz largely disappeared after the aperiodic component was removed, suggesting a change in spectral slope rather than in periodic activity (Figure 5b).

Mean alpha PLV decreased from .553 to .496 (−0.057 [−0.080, −0.035], *q* < .01^**^, all 10 participants). All four regions changed in the same direction (Figure 5c), and 80 of 91 electrode pairs showed a decrease (Figure 5d). Network measures changed correspondingly, with longer characteristic path length and lower clustering coefficient and degree (all 10 participants; Supplementary Appendix 3). A similar change was observed in the beta band but not in the theta band.

### 3.5 Within-person hypotheses and everyday behavior

Within participants, the exponent immediately after a reversal was nearly unchanged from that before it (parietooccipital +0.033 [−0.033, +0.099], *n* = 9; W1). Differences in theta by feedback type were small when all unrewarded trials were pooled (frontal −0.11 dB [−0.68, +0.46], *n* = 10; W2). When unrewarded trials were separated, however, theta following an incorrect choice was higher than theta following a correct choice whose reward was withheld (+0.35 dB [+0.07, +0.59], 7 of 8 participants; family-corrected *q* = .24). Neither preregistered within-person hypothesis was supported in the predicted direction (a lower exponent after reversal; greater theta after unrewarded feedback; models and details in Supplementary Appendix 3).

Informant-reported RBQ-2A sameness scores likewise showed no clear association with task measures (switch bias index ρ = −.25 [−.80, .45], *n* = 12; other measures and the sensitivity scoring are reported in Supplementary Appendix 2). Scores clustered near the floor and internal consistency was low (Cronbach’s α = .38), which limits interpretation.

### 3.6 Validity and signal quality

Between-run reliability of the key measures was high (Figure 5e). ICCs were .945 for the frontal exponent, .903 for frontal alpha peak amplitude, .990 for the occipital (O1, O2) exponent and .818 for occipital alpha peak amplitude; in the preregistered parieto-occipital cluster, they were .984 and .863 (Supplementary Appendix 3). Spectral fits were good (median R^2^ = .98). Because the two runs were recorded consecutively within one session, these values reflect within-session reliability; they are consistent with reports of high test–retest reliability of resting-state spectral measures (Popov et al., 2023).

Spatial distributions were consistent with expectations. Relative alpha power was largest over occipital sites (Figure S1; O1 .364, O2 .354, other channels .13–.24), in line with the posterior dominance of alpha in the eyes-closed state. The channel-wise exponent, for which larger values indicate steeper spectra, was large at frontal sites (1.30–1.45) and small at temporal sites (0.74–1.13), forming an anterior–posterior gradient (Euler et al., 2024). Temporal electrodes, however, are sensitive to muscle activity, which adds high-frequency power and can lead to an underestimated exponent (Goncharova et al., 2003; Whitham et al., 2007). Bad channels were concentrated at P8 (51.6% of runs) and AF4 (41.9%); F7 and T8 were each excluded in 6.5% of runs, and the remaining 10 channels were never excluded.

Of the preregistered feasibility criteria (F1–F7), those for recruitment, practice pass rate, retention and task engagement were met, whereas EEG yield (F3) was met only under the preregistered rule and model fit (F7) could not be assessed with the preregistered index and fell below the threshold with a substitute index (Table 2). Under the manuscript specification with a 60-s minimum per run, EEG yield was 40.0%, below the criterion (80%), which led us to lower the minimum to 30 s. The manuscript specification, however, rejects artifact epochs across more channels than the preregistered rule; under the preregistered cluster-wise rule, yield at 60 s was 12/15 (80.0%), meeting the criterion (Supplementary Appendix 1). Model fit is reported with substitute indices because the preregistered criterion (PSIS-LOO) cannot be computed without a hierarchical Bayesian model.

**Table 2.** Preregistered feasibility outcomes (F1–F7)

|  | Criterion | Result | Threshold | Outcome |
| --- | --- | --- | --- | --- |
| F1 | Recruitment (completed sessions) | 15 | $\geq 10$ | Met |
| F2 | Practice pass | 12/15 (80.0%) | $\geq 70\%$ | Met |
| F3 | EEG yield (parieto-occipital) | Manuscript spec. 60 s 6/15 (40.0%) · 30 s 14/15 (93.3%) · preregistered rule 60 s 12/15 (80.0%) | $\geq 80\%$ | Not met (manuscript spec., 60 s) · met (preregistered rule) |
| F4 | Between-run reliability | ICC .82–.99 | None | — |
| F5 | Retention (withdrawal) | 1/16 (6.2%) | $< 20\%$ | Met |
| F6 | Engagement (median non-response) | 1.4% (range 0.0–19.2%) | $< 10\%$ | Met |
| F7 | Model fit (participants better than random choice) | BIC 7/12 (58.3%) · log-likelihood 12/12 | $\geq 80\%$ (PSIS-LOO) | Preregistered index not computable · BIC below threshold |

*Note.* A completed session is one in which both resting-state runs were recorded after consent. EEG values are based on the manuscript specification (original reference, 200 $\mu\text{V}$ , $R^2 \geq .85$ ). Log-likelihood favors models with more parameters and is therefore less informative. BIC = Bayesian information criterion; ICC = intraclass correlation coefficient; PSIS-LOO = Pareto-smoothed importance sampling leave-one-out cross-validation.

## 4. Discussion

In measuring reversal learning and resting-state EEG together in adults with developmental disabilities in a community setting, we obtained three main findings. First, total errors were comparable across participants, but the type of error split in opposite directions, toward perseveration or regression. Second, exploratory analyses showed that the frontal resting-state aperiodic exponent and alpha peak amplitude were associated with different behavioral outcomes, suggesting potentially distinct behavioral correlates of the two spectral components. Third, during the task, alpha activity and sensor-level alpha synchrony decreased together in almost all participants, and the key resting-state measures were highly reliable within a session.

### 4.1 Responses that do not converge

The switch bias index ranged from −0.965 to +0.909; its mean (+0.510) indicated a net lean toward regressive errors but obscured the two perseverative-dominant participants. When individuals exhibit opposing response tendencies, group-level averages may obscure meaningful heterogeneity. Such heterogeneity may also help contextualize differences in error patterns reported across previous studies (Crawley et al., 2020; D’Cruz et al., 2013). Responses of adults with developmental disabilities may therefore be better described not as a single pattern but together with the direction in which each person leans. In practice, perseverative-dominant and regressive-dominant participants may need different support, and total error scores alone do not distinguish them. For example, perseverative-dominant participants may benefit from advance notice and time to disengage from the old rule, whereas regressive-dominant participants may benefit from consistent confirmation until the new rule is established. These suggestions are hypotheses for future studies and were not tested here.

### 4.2 Resting aperiodic exponent and direction of response

The aperiodic exponent has been interpreted in two main ways. Under the excitation–inhibition interpretation, a steeper spectrum indicates relatively more inhibition whereas a flatter spectrum more excitation (Gao et al., 2017). Under the neural-noise interpretation, a flatter spectrum indicates a state in which responses evoked by external events are less distinguishable from background activity (Voytek et al., 2015). The two views are not mutually exclusive, as increased excitation itself may contribute to neural noise, but they lead to different behavioral predictions. Following the first view, the preregistration expected participants with flatter (excitation-dominant) spectra to switch more readily.

The observed direction was the opposite and is more consistent with the neural-noise view: participants with flatter spectra changed choice less after negative feedback and made more perseverative errors (Figure 4d–f). If a flatter aperiodic spectrum reflects increased neural noise, negative feedback may become less distinguishable from ongoing neural activity and therefore less effective as a signal to switch, potentially leading to repeated selection of a previously rewarded option. Because this interpretation was developed post hoc, and neither neural noise nor feedback sensitivity was directly measured, it requires direct testing in future studies.

Under the same specification, the association was numerically larger in the frontal cluster (ρ = .72) than in the preregistered primary (parieto-occipital) cluster (ρ = .53; Figure 4i); the difference was not tested, and the intervals overlapped widely. If the difference is real, two explanations are plausible. First, prefrontal regions are central to disengaging from a previous choice and moving to a new rule (Uddin, 2021), and frontal electrodes may be more sensitive to activity associated with these processes. Second, the large eyes-closed alpha peak over parietooccipital sites may make it harder to separate the aperiodic component from the periodic one there, even though between-run reliability was high in both clusters.

### 4.3 Alpha peak amplitude and level of performance

Alpha peak amplitude was associated with the level of performance: participants with larger peaks were more accurate and reached the new rule faster (Figure 4g–h), whereas no clear association was observed with the switch bias index or perseverative errors. Alpha has been interpreted as a rhythm that gates information flow by inhibiting task-irrelevant processing (Jensen & Mazaheri, 2010); from this perspective, individuals with a larger resting alpha peak may process task-relevant information more efficiently. A similar dissociation has been reported in visuospatial attention, where the periodic alpha component was linked to accuracy and the aperiodic component to response speed (Lu et al., 2025).

The respective associations remained in partial correlations controlling for the other EEG measure (Figure 4j), but the intervals for the differences between correlations (Δρ) included zero. Therefore, the dissociation cannot be established from these data alone. A direct test would manipulate behavior linked to one measure: lowering the reward probability to weaken error signals would be expected to strengthen the link between the exponent and lose-shift, whereas degrading stimuli to make the task harder would be expected to strengthen the link between alpha peak amplitude and accuracy.

### 4.4 From rest to task

When the task began, alpha activity and alpha synchrony between electrode groups decreased together in almost all participants. Because rest was recorded with eyes closed and the task was performed with eyes open, part or all of this decrease may reflect eye opening alone. Alpha suppression is also a typical response to task engagement; if alpha inhibits task-irrelevant processing (Jensen & Mazaheri, 2010), the decrease may partly reflect a release, in the service of task processing, of the inhibition that was widespread at rest. The reduced sensor-level synchrony and network changes (longer characteristic path length, lower clustering coefficient) are compatible with this interpretation, although scalp PLV cannot show whether cortical regions became less coupled. The decrease was observed in every electrode grouping, with widely overlapping intervals (Figure 5c).

These results are also informative about the nature of the individual differences. The way the task changed brain state was common to most participants, whereas the direction of response after reversal differed between individuals; the differences lay less in the capacity to engage with the task than in how feedback was used once engaged. The finding that signals recorded with a mobile device in the field changed in the expected direction also suggests that the measurement captured real state changes. Separating task engagement from eye opening would require an eyes-open resting condition.

### 4.5 Within-person change and everyday behavior

Within participants, the exponent hardly changed immediately after reversals (W1), and theta differences by feedback type were small when unrewarded trials were pooled (W2). This is consistent with the possibility that the exponent reflects relatively stable individual differences over the timescale examined: in the reversal-window model, most of the variance lay between participants (about 22 times the variance between window pairs within participants at parieto-occipital sites), and between-run reliability was high. Taken together, these findings suggest that resting-state differences were more closely associated with individual response tendencies than the reversal-related changes in exponent examined here.

Unrewarded trials comprise two kinds of events: incorrect choices, and correct choices whose reward was withheld under the probabilistic (80%) schedule. The outcome on screen (X) is the same, but only in the former was the choice actually incorrect. Theta was higher in the former (+0.35 dB [+0.07, +0.59], 7 of 8 participants; Section 3.5), suggesting that theta may respond to whether one’s choice was incorrect rather than to the absence of reward. This may also explain the small difference in the preregistered pooled comparison (W2). Because this result is based on eight participants with a corrected *q* of .24, it requires confirmation with a prespecified contrast in future studies.

Informant-reported everyday repetitive behavior showed no clear association with task measures. Because this scale has not been validated in adults like our participants (Brett et al., 2024) and scores clustered near the floor, narrowing the range of individual differences, the measure was likely unable to reveal an association rather than showing that none exists; studies using validated instruments have reported such associations in young autistic children (Saniee et al., 2026).

### 4.6 Validity and field application

Although the device was consumer-grade, the key measures showed high reliability. ICCs of .82–.99 fall in the good-to-excellent range by common standards (Koo & Li, 2016), consistent with high test–retest reliability of resting spectral measures with research-grade systems (Popov et al., 2023); spectral fits had a median R^2^ of .98, and spatial distributions were physiologically plausible. Because the two runs were consecutive within one session, reliability across days remains untested.

Feasibility was evaluated against the preregistered criteria (Table 2). Fifteen participants completed the session, and one (6.2%) withdrew; the practice pass rate (80.0%) exceeded the criterion (70%). The median nonresponse rate was 1.4%, indicating that participants stayed engaged throughout the 20-min session. Under the preregistered rule, EEG yield was exactly at the criterion (80.0%); data loss arose mainly from poor electrode–skin contact at P8 and AF4 (bad-channel rates of 51.6% and 41.9% under the manuscript specification). Thus, the measurement was feasible for most participants at a community day-service center within 20 min and without sedation, with the remaining data loss narrowed down to specific electrodes. Mobile EEG has been proposed as a means of lowering the barriers to data collection in developmental conditions (Lau-Zhu et al., 2019); our results support this possibility in a sample that included adults with intellectual disability. Recording with low-cost equipment at places participants already attend offers a practical route for including people who have been left out of research in the evidence about them.

### 4.7 Limitations and future research

With primary samples of 9–15 participants, intervals were wide, and only two participants were perseverative-dominant. The preregistration targeted autistic adults, but the sample consisted of adults with registered developmental disabilities (four with autism), and without a comparison group we cannot determine whether the individual differences are specific to this population. The sample was too small to control for medication and age. Rest (eyes closed) and task (eyes open) conditions differed, the device lacked midline electrodes and had a fixed reference, and the switch bias index overlaps structurally with the overall switch rate (a property stated in the preregistration from simulations of the fixed schedule; ρ = .95 in the data, *n* = 9; Supplementary Appendix 1). Everyday behavior was measured with a single questionnaire not validated in this population, and interpretations of results opposite to the prediction are post hoc. Finally, the study was conducted without formal IRB review, so the results should be read as preliminary observations.

Future studies should recruit larger samples across multiple centers to narrow the intervals and secure enough people of both response types, increase the number of trials and reversals, add eyes-open rest and validated adult measures of everyday behavior, and manipulate the strength of feedback signals to test the neural-noise interpretation directly.

### 4.8 Conclusion

Using mobile EEG in a community setting, we measured behavior and resting-state EEG together in adults with developmental disabilities, including those with intellectual disability. After reversals, individual responses diverged toward opposite poles, and in exploratory analyses this divergence was associated with the frontal resting-state aperiodic exponent. These results provide preliminary evidence that field-recorded EEG can capture individual differences in how people respond to change, and they offer effect sizes and design requirements for future studies. If episodes of challenging behavior coincide with changes in rules and environments, measuring how each person responds to change may provide a starting point for designing announcements and guidance tailored to each individual. Extending such evidence to people with intellectual disability is a shared task for research on health and well-being (SDG 3) and reduced inequalities (SDG 10).

## Declarations

### Ethics statement

This study was conducted in accordance with the principles of the Declaration of Helsinki. The protocol and consent procedure were reviewed in a mock review within the Social Engagement Fund program of the Institute for Global Engagement & Empowerment, Yonsei University; this was not a formal review by an institutional review board. The study was conducted under a letter of cooperation from the participating center.

### Consent to participate

All participants were adults and, after the study was explained, signed the consent form themselves. People under adult guardianship were not eligible; none of those approached was under guardianship.

### Data availability

Because the data were collected without institutional review board approval, raw data are neither made public nor shared with third parties. Participant-level summary values used in the analyses are provided in Tables S2 and S3.

### Code availability

Analysis code is available in a public repository (https://github.com/JeongNeuro/reversal-learningeeg); raw data are not included. Behavioral, resting spectral and connectivity measures and all figures are reproduced from the raw data with this code. Figures 1, 4 and 5 were edited for layout from the code output, and the scatter plots and fits in Figure 4d–h and the matrix in Figure 5d were redrawn from the same values; no numbers changed. The original script for the W1 window exponents was not retained; the repository rebuilds windows from the preregistered specification, and the conclusions on sign and intervals in all four clusters are the same in both versions. The PsychoPy task script, which classifies perseverative and regressive errors, and the fixed-seed trial schedule are also released.

### Author contributions

Yunjeong Jeong: conceptualization, methodology, investigation, formal analysis, data curation, visualization, writing – original draft, writing – review and editing, project administration. Suho Hyun: conceptualization, investigation, writing – review and editing. Both authors approved the final manuscript.

### Use of AI tools

A large language model (Anthropic Claude) was used to write and debug analysis code (preprocessing, statistics, figure scripts), to implement the PsychoPy task program, and to edit the wording of the manuscript and check numbers and notation. All decisions on study design, analytic methods and criteria, interpretation of results and the final content were made by the first author, who reviewed all model-written code and verified its output before use. The authors take full responsibility for the content.

## Supplementary material

*Responses to unannounced rule changes in adults with developmental disabilities: A community-based pilot study with consumer-grade EEG*

## Appendix 1. Analysis samples and deviations from the preregistration

**Table S1.**
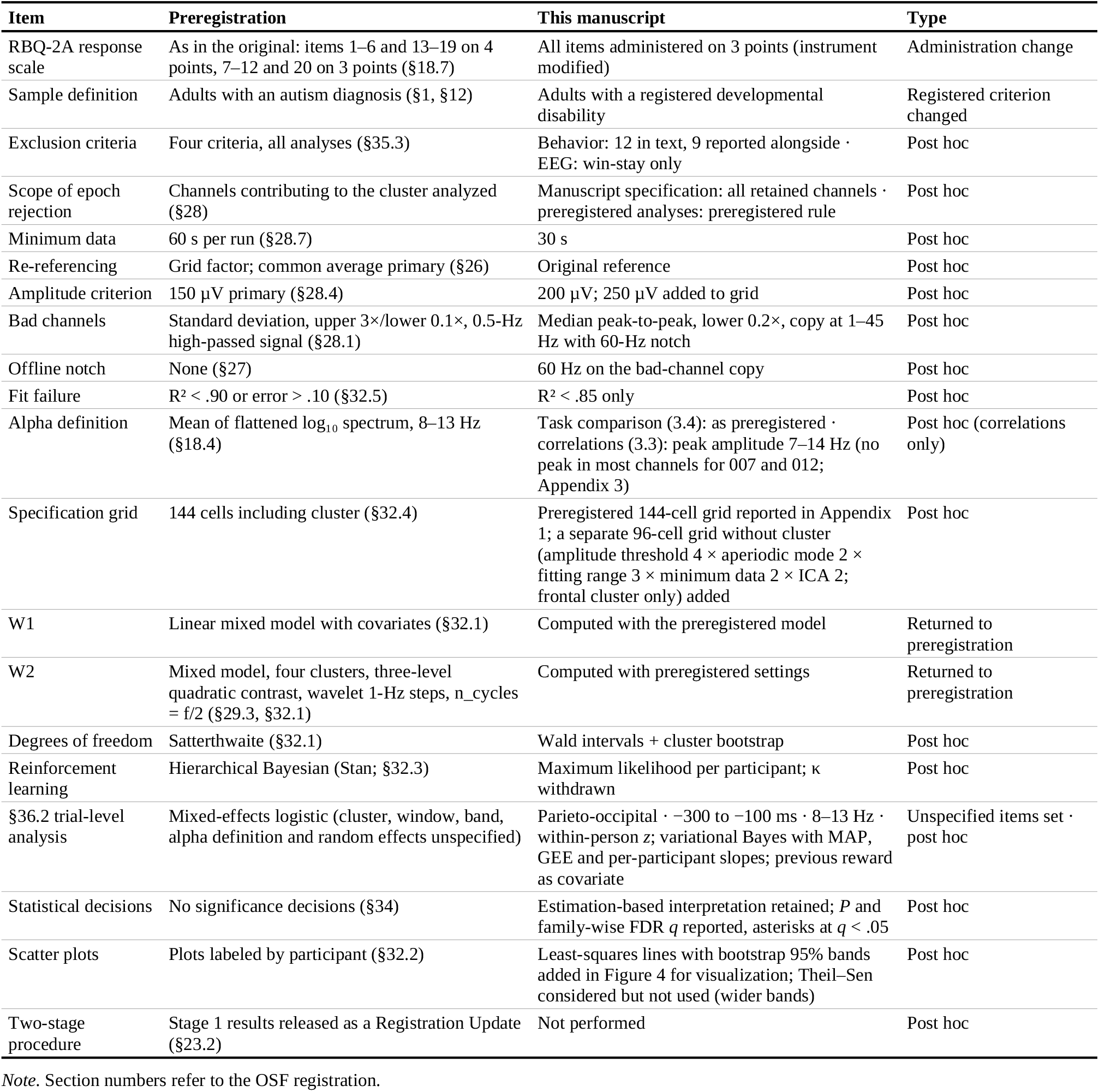
Deviations from the preregistration.

### Robustness

The association between the frontal exponent and lose-shift was largely insensitive to analytic choices. Leave-one-out estimates ranged from .738 to .838 (.790 when 004 was left out); reinstating 009, who had been excluded by the win-stay criterion (*n* = 10; 007 remained excluded by the reaction-time criterion), yielded .766 [.27, .94]; and applying none of the exclusion criteria (*n* = 12) yielded .655 [.16, .87]. The association with the switch bias index ranged from .595 to .833 in leave-one-out analyses (.667 when 004 was left out) and was .612 [−.22, .91] at *n* = 10, but it weakened to .112 [−.52, .73] at *n* = 12.

Two specification grids are reported. The 96-cell grid listed in Table S1 is a separate post hoc grid that drops cluster as a factor and varies the amplitude threshold (4), aperiodic mode (2), fitting range (3), minimum data per run (2) and ICA (2) for the frontal cluster only. Of the 144 specifications in the preregistered grid (§32.4; reference 2 × amplitude threshold 3 × fitting range 3 × aperiodic mode 2 × cluster 4, with a 60-s minimum per run), all 41 specifications that met the preregistered minimum of eight participants yielded positive estimates (median +0.50, range +0.26 to +0.78). Across all 144 specifications, 119 (0.826) matched the sign of the preregistered specification, falling short of the preregistered sign-stability criterion (0.90); all 25 specifications with the opposite sign were below the minimum sample size (§35.9). With a 30-s minimum, the corresponding proportions were 0.896 across all 144 specifications and 1.000 across the 76 specifications that met the minimum.

Age was not associated with the frontal exponent (ρ = −.301 [−.75, .28], *n* = 15) but was associated with the occipital (O1, O2) exponent (ρ = −.592 [−.88, −.11], *n* = 14).

The result for the primary hypothesis under the preregistered specification (common average reference, 150 µV, 2–40 Hz, fixed mode, parieto-occipital cluster, at least 60 s per run; ρ = +.69, *n* = 8: 001–006, 012, 014) is reported in Section 3.3. Lowering the minimum to 30 s per run yielded ρ = +.67 [−.11, .98] (*n* = 9, *P* = .061). Behavioral values for the 9 participants who remained after all preregistered exclusion criteria were applied (001–006, 012, 014, 016) are given in Table S3. The three participants excluded were 007 (24.2% of trials with reaction times < 150 ms), 008 (win-stay .595) and 009 (win-stay .538); 006 approached the 20% criterion with 19.2% non-responses, and 009 had exactly the minimum of 96 valid trials. In this sample, the switch bias index was correlated with the switch rate at ρ = .95 (Pearson *r* = .94). The mean switch bias index leaned toward regressive errors (+0.39), but its interval included zero ([−0.02, 0.72]); 7 of the 9 values were positive.

### Deviations from the preregistration

The criteria for bad-channel detection, amplitude-based rejection and minimum data per run were adjusted after the data had been inspected. With a 60-s minimum per run, only 10 of 31 runs remained, and the minimum was therefore lowered to 30 s (parieto-occipital yield, 6/15 at 60 s and 14/15 at 30 s; Table 2). This yield reflects the manuscript specification, in which artifact epochs are rejected on the basis of all retained channels. When epochs were instead judged only on the channels contributing to each cluster, as specified in §28, 12/15 participants met the minimum at 60 s; the preregistered analyses (the preregistered specification in Section 3.3 and the specification grid) therefore used this clusterwise rule. Although the preregistration specified the common average reference as primary, the 14 channels are unevenly distributed over the scalp; the main text therefore reports results with the original reference and presents common-average results alongside (post hoc decision). The two-stage procedure of §23 was not performed. The alpha peak amplitude (7–14 Hz) used in the correlational analyses (Section 3.3) differs from the preregistered definition of periodic alpha (§18.4), whereas the rest–task comparison (Section 3.4) followed the preregistered definition. The topographies in Figure 5a are based on channel-wise relative power and are therefore derived differently from the cluster values. All deviations are listed in Table S1. In a sensitivity analysis with independent component analysis, the direction of the primary associations was unchanged (e.g., exponent × lose-shift, ρ = .803 → .739).

### Participant-level values

For this sample, the preregistration specified participant-level values, rather than correlation coefficients, as the primary output. A correlation coefficient assumes a single monotonic association across all participants, and opposing pathways within the sample may cancel each other in such a summary.

**Table S2.**
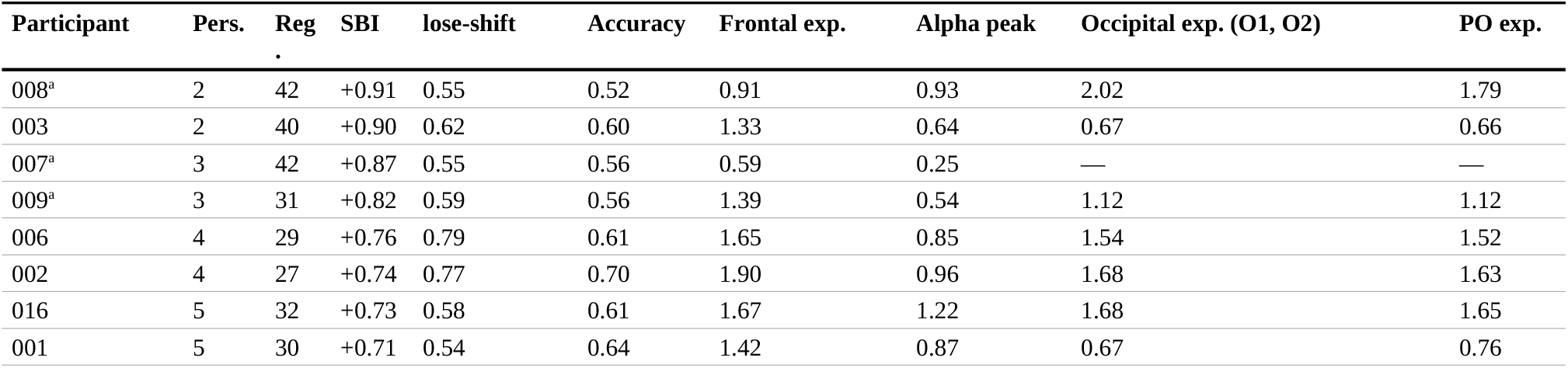
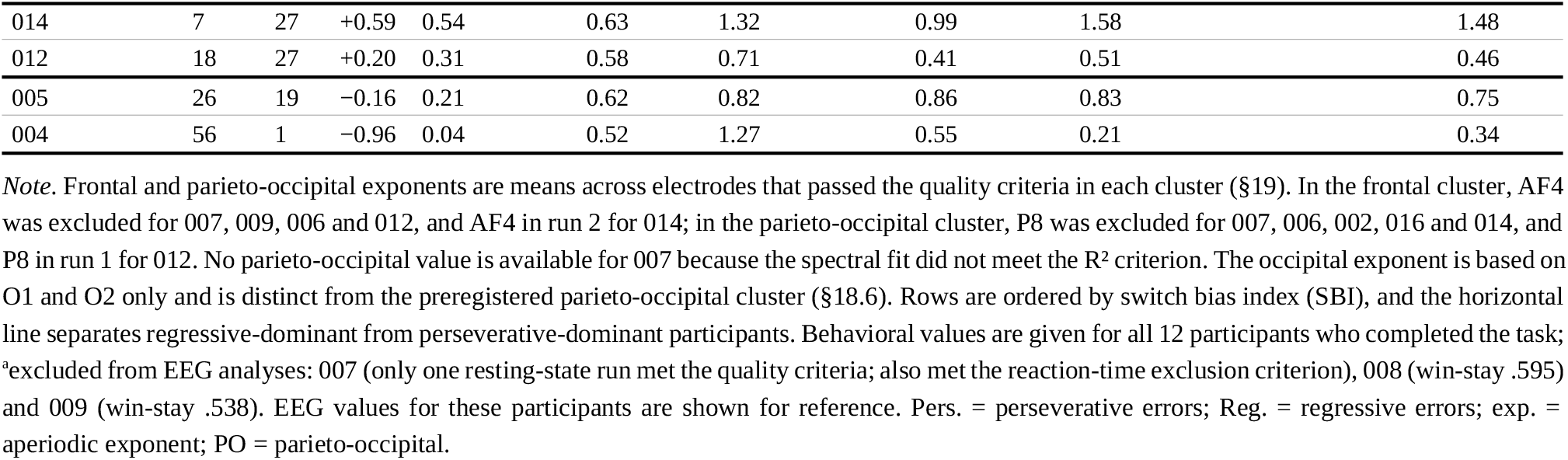
Participant-level values, ordered by switch bias index.

**Table S3.** Behavioral measures in the sample meeting all four preregistered criteria (§35.3; *n* = 9)

| Participant | Perseverative errors | Regressive errors | SBI | lose-shift | win-stay | Accuracy | Switch rate | Trials to recover |
| --- | --- | --- | --- | --- | --- | --- | --- | --- |
| 001 | 5 | 30 | +0.714 | .538 | .815 | .642 | .378 | 3.80 |
| 002 | 4 | 27 | +0.742 | .770 | .966 | .700 | .412 | 6.80 |
| 003 | 2 | 40 | +0.905 | .623 | .653 | .597 | .508 | 8.00 |
| 004 | 56 | 1 | -0.965 | .043 | .980 | .517 | .034 | 11.20 |
| 005 | 26 | 19 | -0.156 | .210 | .965 | .625 | .126 | 8.40 |
| 006 | 4 | 29 | +0.758 | .792 | .744 | .608 | .552 | 10.20 |
| 012 | 18 | 27 | +0.200 | .306 | .833 | .577 | .245 | 12.40 |
| 014 | 7 | 27 | +0.588 | .538 | .778 | .633 | .395 | 6.40 |
| 016 | 5 | 32 | +0.730 | .582 | .708 | .612 | .461 | 5.40 |
| **Mean** | 14.1 | 25.8 | +0.391 | .489 | .827 | .612 | .346 | 8.07 |
| SD | 17.6 | 10.8 | 0.608 | .253 | .120 | .050 | .175 | 2.81 |
| 95% CI | [5.1, 26.1] | [18.6, 31.7] | [-0.02, +0.72] | [.33, .64] | [.75, .90] | [.58, .64] | [.23, .45] | [6.4, 9.8] |

*Note.* Excluded participants: 007 (only one resting-state run met the quality criteria; 24.2% of trials with reaction times < 150 ms, above the preregistered limit of 20%), 008 (win-stay .595 < .60) and 009 (win-stay .538 < .60). The 95% confidence intervals of the means are based on 10,000 participant resamples (seed 1). SBI = switch bias index (+1, predominantly regressive; -1, predominantly perseverative).

## Appendix 2. Supplementary behavioral analyses

### Reinforcement learning model

The preregistered model included *η*_rew_, *η*_unrew_, *β* and *κ*, with *κ* registered as a co-primary outcome, and specified hierarchical Bayesian estimation; the model was instead estimated by maximum likelihood for each participant. In simulated data, the recovery correlation for *κ* (+.924) exceeded the preregistered criterion (0.60). In the observed data, however, learning rates converged to their bounds and inverse temperatures diverged, so the parameters could not be identified. Because 120 trials and five reversals appear insufficient to identify this model, *κ* was withdrawn as a co-primary outcome and replaced by the lose-shift rate (Table S1). The model fit better than random choice for 7 of 12 participants (58.3%) by the Bayesian information criterion (BIC), below the preregistered criterion of 80% (F7).

### RBQ-2A and task measures

Associations between informant-reported RBQ-2A sameness scores (preregistered primary scoring: the 13-item structure of Barrett et al., 2018) and behavioral measures all had wide intervals (switch bias index, ρ = −.25 [−.80, .45]; perseverative errors, +.20 [−.51, .78]; regressive errors, −.24 [−.81, .41]; lose-shift, −.06 [−.70, .62]; all *n* = 12). The preregistered sensitivity scoring (the 7-item structure of Barrett et al., 2015) yielded nearly identical values. Scores lay near the floor (13–18 of a possible 13–39), and internal consistency was low (Cronbach’s α = .38, 14 respondents). Because items originally rated on a 4-point scale were administered on a 3-point scale, scores are not on the original metric (Table S1).

## Appendix 3. Supplementary EEG analyses

**Figure S1.**
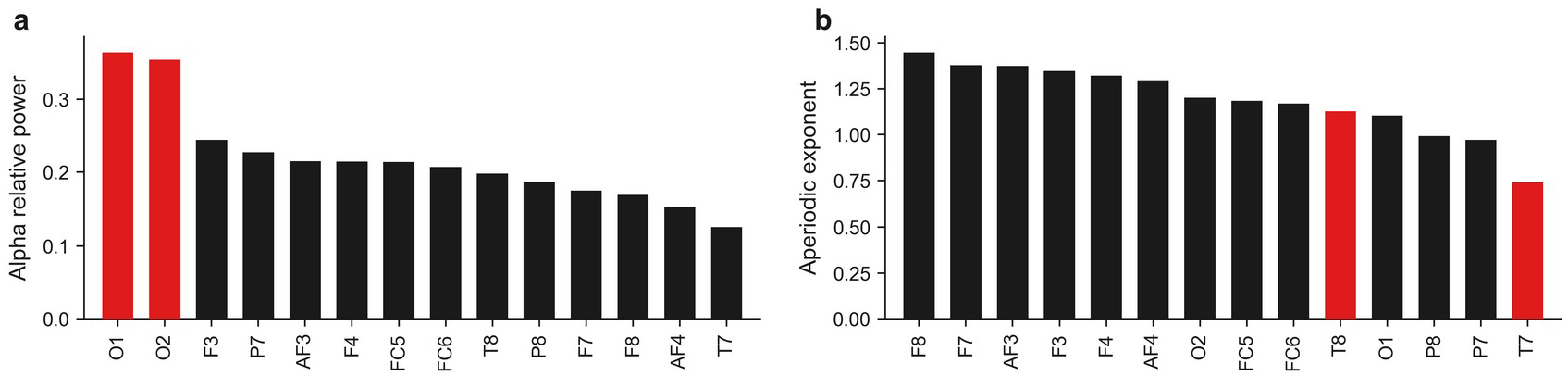
Group means (*n* = 15) of channel relative alpha power (a) and aperiodic exponent (b). Red marks the two occipital electrodes in a and the two temporal electrodes in b

### Validity and stability

Under the preregistered specification (common average reference, 150 µV, R^2^ ≥ .90 and error ≤ .10, cluster-wise epoch rejection), the parieto-occipital yield was 12/15 (80.0%) with a 60-s minimum, and the between-run ICC for the exponent was .959 [.88, .99]. Values under the manuscript specification are given in Section 3.6 and Table 2; under that specification, the parieto-occipital (O1, O2, P7, P8) ICCs were .984 [.96, .99] for the exponent and .863 [.44, .98] for alpha peak amplitude (both *n* = 11). The occipital measure (O1, O2) is based on two electrodes only and is distinct from the parieto-occipital cluster.

In an exploratory comparison of the two runs (§36.1), the exponent was lower in the second run at parieto-occipital (−0.058 [−0.100, −0.017]) and frontal sites (−0.080 [−0.139, −0.021]; both *n* = 11). Because the procedure and interval between runs were not fixed, this design cannot distinguish adaptation from measurement instability.

The median clock offset drift was +0.05 ms, and the median proportion of rejected epochs per run was 35.0% (range 0.0–77.5%; §28). A 7–14 Hz alpha peak was fitted in 168 of 196 channels (85.7%) but rarely in 007 (1/12) and 012 (3/13), whose peak measures should therefore be considered unstable. Stimulus A (blue circle) was chosen on 64.1% of trials on average (range 50–98%); because stimulus A was always the correct option in the first segment, this proportion is reported descriptively only (§7.1, §36.7).

### Spectral and network changes during the task

Values in parentheses are paired differences (task − rest) with bootstrap 95% intervals. In the frontal cluster (AF3, AF4, F3, F4), periodic alpha after removal of the aperiodic component (the preregistered definition, §18.4) decreased from .477 to .190 (−0.287 [−0.407, −0.162], *P* = .016, *q* < .05^*^, 7 of 8 participants); the decrease in relative alpha power yielded *P* = .006. Alpha peak amplitude estimated with specparam decreased from 1.160 to 0.512 in the occipital cluster (−0.648 [−0.836, −0.444], all 8 participants) and from 0.806 to 0.387 in the frontal cluster (−0.419 [−0.614, −0.236], 6 of 7). Frontal relative delta power increased from .157 to .200 (*P* = .014), whereas intervals for theta (+0.021 [−0.005, 0.049]), beta (−0.001 [−0.022, 0.025]) and gamma (+0.012 [−0.010, 0.041]) included zero. Peak frequency shifted by −0.75 Hz frontally (9.21 to 8.46 Hz; [−1.55, −0.03]) and by +0.63 Hz occipitally (8.92 to 9.55 Hz; [−0.17, 1.64]); given the opposite directions and wide intervals, the size and direction of any frequency shift cannot be determined from these data.

The decrease in mean alpha PLV yielded *P* = .002. Regional decreases were frontal −0.068 [−0.097, −0.044], frontoposterior −0.059 [−0.092, −0.030], posterior −0.055 [−0.081, −0.031] and central −0.048 [−0.074, −0.024] (central *q* < .05^*^, others *q* < .01^**^); because the intervals overlap widely, no ordering of regions can be inferred. Characteristic path length increased from 1.91 to 2.11 (+0.195 [0.123, 0.271]), the clustering coefficient decreased from .618 to .581 (−0.038 [−0.058, −0.020]), and the number of suprathreshold connections decreased from 6.17 to 3.75 (−2.42 [−3.42, −1.43]; all 10 participants).

### Within-person hypothesis W1

The exponent in the eight trials after each reversal was compared with that in the eight trials before it, using the preregistered linear mixed model (window, reward density, window length and epoch count; random intercepts for participant and participant × pair). At parieto-occipital sites the exponent was +0.033 [−0.033, 0.099] higher in post-reversal windows (9 participants, 36 pairs, *P* = .32); other clusters gave frontal +0.027, temporo-parietal +0.044 and global +0.036. Between-participant variance was approximately 22 times the variance between window pairs within participants. Because the reward schedule was fixed, reward density did not vary between participants and therefore could not control for imbalance. The median window length was 22.8 s across all cluster rows (23.2 s across distinct windows; range 18.0– 37.8 s), corresponding to a median of 10–11 artifact-free 2-s epochs. The 60-s minimum of §28.7 applies to resting-state runs, not to these eight-trial task windows; the floor applied here was five artifact-free epochs (10 s) per window and per cluster, counted after the cluster-wise amplitude rejection of §28, so the number of surviving epochs could differ between clusters for the same window. No window survived for 001 or 007, and the manuscript specification yielded nearly the same result (parieto-occipital +0.032 [−0.031, 0.094]).

### Within-person hypothesis W2

Under the preregistered specification, feedback-locked theta (200–500 ms, 4–8 Hz) differed between unrewarded and rewarded trials by −0.11 dB [−0.68, 0.46] frontally (*n* = 10), −0.40 [−1.09, 0.28] parieto-occipitally (*n* = 12), +0.06 [−0.49, 0.62] temporo-parietally (*n* = 10) and −0.09 [−1.08, 0.91] globally (*n* = 9); a trial-level random-slope model gave the same direction (frontal −0.17 [−0.78, 0.44], 922 trials). Sample sizes differ across clusters because participants were included only if at least 10 epochs per condition remained in a given cluster (§29). Because unrewarded trials comprise both incorrect choices and correct choices whose reward was withheld, the preregistered three-level contrast was also computed (*n* = 8): relative to rewarded-correct trials, frontal theta was −0.59 dB [−1.05, −0.07] for unrewarded-correct and −0.24 dB [−0.72, 0.28] for unrewarded-incorrect trials (family-corrected *q* = .24 and .66, respectively).

### Multivariate correspondence (Mantel)

In the preregistered multivariate correspondence analysis (Spearman correlations between participant-by-participant distance matrices for the behavioral, model and EEG blocks; 10,000 label permutations), no correspondence between behavior and EEG was observed (frontal ρ = −.06, parieto-occipital −.05, both *n* = 9). The model block was not interpreted because its parameters could not be identified, and at this sample size the Mantel test cannot establish the absence of a correspondence.

### Pre-stimulus alpha and switching

With the preregistered trial-level mixed-effects logistic model (switch ∼ pre-stimulus alpha + participant random intercepts and slopes; 9 participants, 994 trials), we tested whether alpha power (8–13 Hz, within-person *z*) at −300 to −100 ms before the stimulus predicted switching on the next choice. In the primary parieto-occipital cluster, the odds ratio per standard deviation of alpha was 0.95 [0.83, 1.10], with identical values across four estimation methods (variational Bayes, MAP, GEE and mean per-participant slopes); when excluded participants were included (*n* = 12), the odds ratio was 1.04 [0.93, 1.18]. The three non-primary clusters showed larger effects in the same direction (e.g., frontal, 0.83 [0.72, 0.96]); because clusters and time windows were not prespecified, however, these results remain exploratory and require confirmation.

**Table S4.**
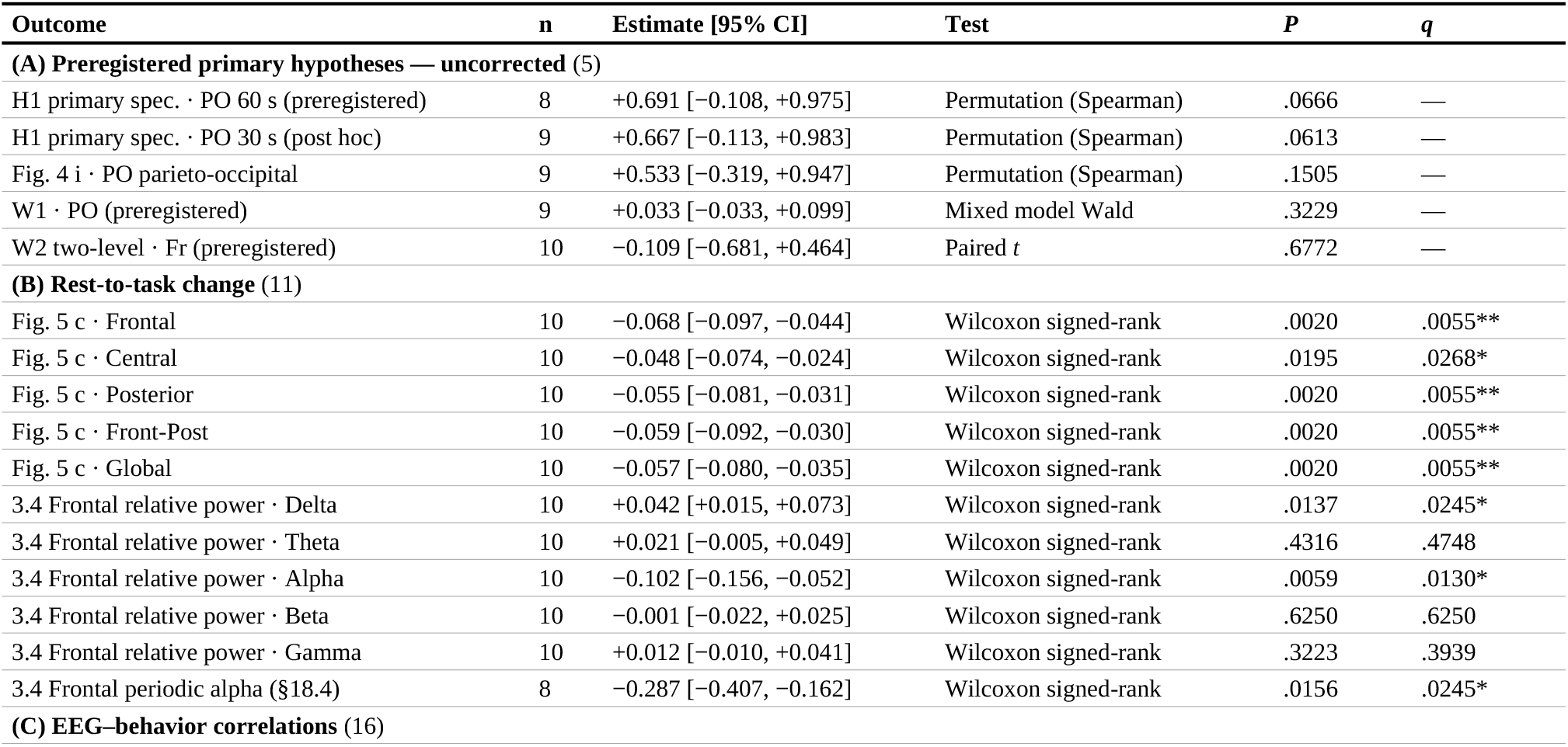
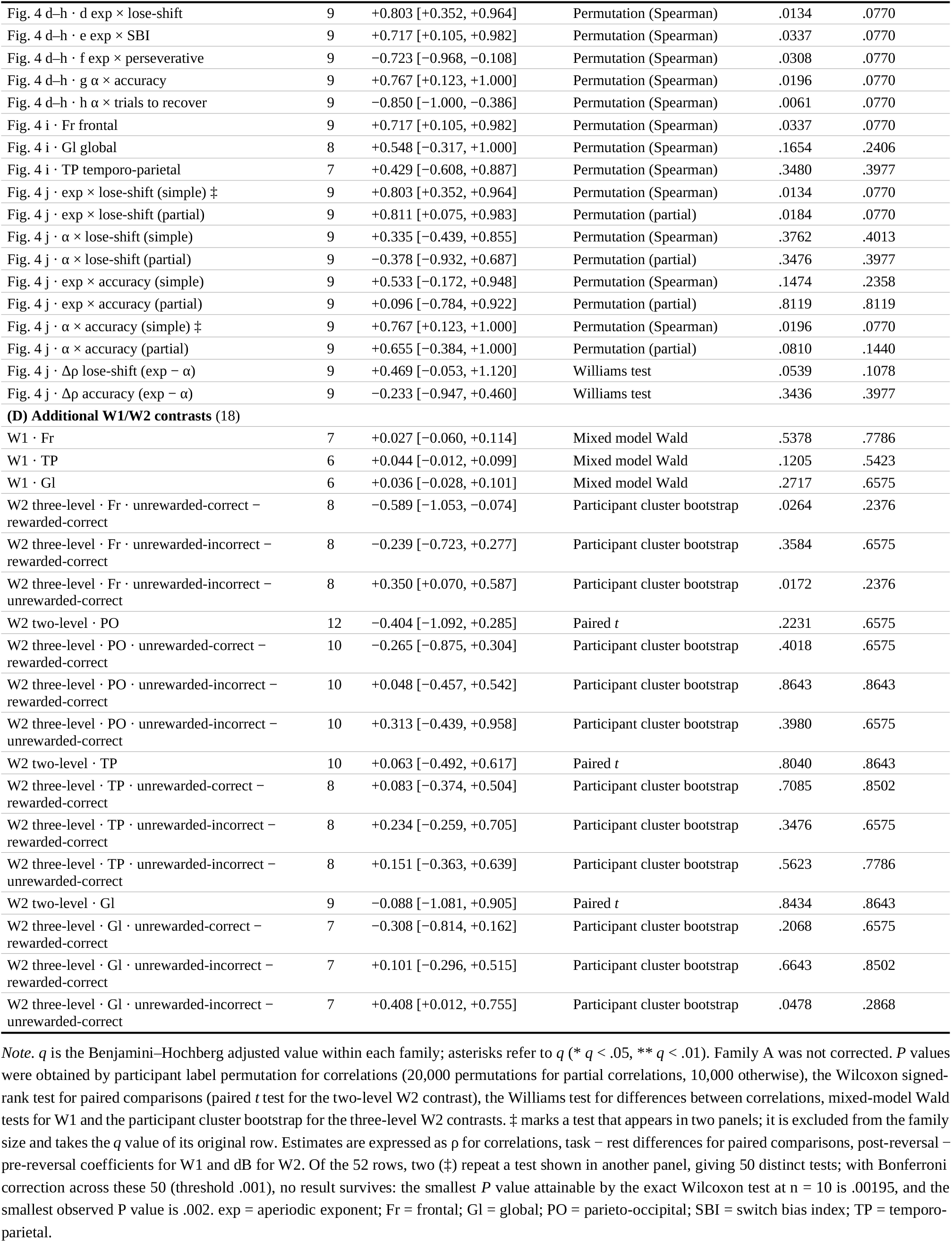
Estimates, tests, *P* values and family-wise *q* values by outcome.

